# Milking it for all it’s worth: The effects of environmental enrichment on maternal nurturance, lactation quality, and offspring social behavior

**DOI:** 10.1101/2021.12.13.472509

**Authors:** Holly DeRosa, Salvatore G. Caradonna, Hieu Tran, Jordan Marrocco, Amanda C. Kentner

**Affiliations:** School of Arts & Sciences, Health Psychology Program, Massachusetts College of Pharmacy and Health Sciences, Boston Massachusetts, United States 02115; Laboratory of Neuroendocrinology, The Rockefeller University, 1230 York Ave, New York, NY 10065; Department of Biology, Touro University, 227 W 60th St, New York, NY 10023

**Author notes:** Author Contributions, H.D., S.G.C., H.T., and A.C.K., ran the experiments; H.D., S.G.C., J.M., & A.C.K. analyzed and interpreted the data; H.D. and A.C.K. wrote the manuscript; A.C.K., designed and supervised the study. Corresponding author: Amanda Kentner, Office.

**Keywords:** Environmental enrichment, animal housing, animal welfare, lactation, nutrition, social behavior, prenatal experience, postnatal experience, press posture, RNA-sequencing, transcriptome, microbiome, milk, breastfeeding, cannabinoid receptors, translational research

## Abstract

Breastfeeding confers robust benefits to offspring development in terms of growth, immunity, and neurophysiology. Similarly, improving environmental complexity (i.e., environmental enrichment; EE) contributes developmental advantages to both humans and laboratory animal models. However, the impact of environmental context on maternal care and milk quality has not been thoroughly evaluated, nor are the biological underpinnings of EE on offspring development understood. Here, Sprague-Dawley rats were housed and bred in either EE or standard-housed (SD) conditions. EE dams gave birth to a larger number of pups, and litters were standardized and cross-fostered across groups on postnatal day (P)1. Maternal milk samples were then collected on P1 (transitional milk phase) and P10 (mature milk phase) for analysis. While EE dams spent less time nursing, postnatal enrichment exposure was associated with heavier offspring bodyweights. Milk from EE mothers had increased triglyceride levels, a greater microbiome diversity, and a significantly higher abundance of bacterial families related to bodyweight and energy metabolism. These differences reflected comparable transcriptomic changes at the genome-wide level. In addition to changes in lactational quality, we observed elevated levels of cannabinoid receptor 1 in the hypothalamus of EE dams, and sex- and time-dependent effects of EE on offspring social behavior. Together, these results underscore the multidimensional impact of the combined neonatal and maternal environments on offspring development and maternal health. Moreover, they highlight potential deficiencies in the use of “gold standard” laboratory housing in the attempt to design translationally relevant animal models in biomedical research.

**Significance Statement:** Maternal care quality is different between environmentally enriched (EE) and standard laboratory housed (SD) dams. SD rat dams spend more time nursing their young. This may result in overfeeding which can program offspring neural stress responses. Alternatively, metabolic differences in milk may affect neurodevelopmental outcomes, which are different between EE and SD animals. To test this, we evaluated milk and offspring behavior. Milk from EE dams had elevated triglyceride levels and microbiome diversity. EE offspring had heavier body weights and increased social behavior which was lost with crossfostering into SD housing. These data identify potential deficiencies in the quality of “gold standard” laboratory housing and its impact on the welfare and design of translationally relevant animal models in biomedical research.

## Introduction

The maternal-infant interaction of breastfeeding is a critical component in the neurodevelopment of offspring. Breastmilk promotes increased white and grey matter volume, improved cortical thickness (Isaacs et al., 2008; Luby et al., 2016) and neuromuscular development (Grace et al., 2017). Milk may facilitate development of the central nervous system by directly influencing genes related to neural growth and maturation. For example, microRNAs in mouse milk target proteins in the offspring small intestine related to neuron proliferation, differentiation, and survival (Huff et al., 2020). While this evidence implicates milk in brain development, the role of milk in offspring behavior needs further elucidation.

The profile of breastmilk can be modulated by the maternal environment, and several studies have revealed the pervasive effects of stress on milk production and quality. For example, mothers who have been exposed to natural disasters often report a reduction or sometimes complete loss of lactation (Adhisivam et al., 2006; DeYoung et al., 2018). Non-disastrous contexts such as psychosocial stress is also negatively associated with maternal milk fat and energy content (Ziomkiewicz et al., 2021) and mothers who reported higher levels of perceived stress exhibited significantly lower levels of milk immunoglobulin A (Moirasgenti et al., 2019; Groer et al., 2004). Likewise, environmental stressors correlate with altered maternal milk quality in laboratory animals. For example, restraint stress has been shown to reduce milk protein in lactating mice (Chiba et al., 2019), and other stressors such as social and heat stress have been shown to negatively affect lactation (Murgatroyd et al., 2015) and milk yield (Haldar & Bade, 1981) respectively. Such evidence underscores how the maternal environment may contribute to alterations in the nutritional profile of milk which may have subsequent consequences on progeny development.

The enhancement of environmental complexity (i.e., environmental enrichment or EE) is employed in healthy human populations to promote cognitive plasticity (Baroncelli et al., 2010; Kentner et al., 2019a; Kentner et al., 2019b; Tooley et al., 2021). Additionally, EE housing reduces stress and stereotypy and promotes species typical behaviors in the animal laboratory. This enhanced housing condition has been shown to affect the display of rodent maternal care behaviors (Welberg et al., 2006; Connors et al., 2015; Cancedda et al., 2004; Mann & Gervais, 2011; Strzelewicz et al., 2019; Sale et al., 2004) which are central to the development of effective stress regulation and health of the offspring (Meaney & Szyf, 2005; Meaney, 2010; Buschdorf & Meaney, 2011). Research has supported the notion that EE dams are more efficient mothers compared to their standard-housed (SD) counterparts. For example, while rats housed in EE spent less time on their nest compared to SD dams, both groups licked and groomed their pups at a similar (Welberg et al., 2006; Connors et al., 2015; Strzelewicz et al., 2019) or even higher frequency (Sale et al., 2004), although these findings are not always consistent (Cancedda et al., 2004; Rosenfeld & Weller, 2012). Additionally, EE dams spend less time in a passive nursing posture compared to SD dams and demonstrate higher levels of the more effective active, or high arched back nursing posture (Connors et al., 2015; Strzelewicz et al., 2019). In another study, rat dams housed in cages with a loft that afforded an opportunity to periodically leave their pups also exhibited lower levels of passive nursing (Ratuski & Weary, 2021). This suggests that time away from the pups promotes an efficient maternal care style upon returning to the nest. This is congruent with what is typically observed under naturalistic conditions, where wild rat dams will leave their nests for extended periods to forage and defend their territory (Hughes et al., 1978; Grota & Ader, 1969).

Given that SD dams housed in the classic “gold standard” laboratory housing condition spend more time on the nest suggests they may overfeed their offspring. Alternatively, there may be metabolic differences in milk quality where SD offspring require more nourishment, necessitating longer nursing periods. Indeed, dams in cages with reduced opportunities to leave their litters spend more time in a “press posture” position with the ventral surface of their body pressed against the cage, hiding their teats from their pups (Gaskill & Pritchett-Corning, 2015; Cramer et al., 1990). Overall, these considerations have important implications for laboratory animal health and stress regulation, which may impact the translational relevance of our animal models. Therefore, in the current study we explored the effects of EE versus SD housing on rodent maternal care, maternal milk quality, and offspring body weight and social behavior outcomes in male and female rats.

## Materials and Methods

### Animals and housing

Sprague-Dawley rats (Charles River, Wilmington, MA) were maintained at 20°C on a 12 h light/dark cycle (0700-1900 light) with *ad libitum* access to food and water. **Figure 1A** outlines the experimental procedures followed in this study. Female animals were pair-housed in one of two conditions: environmental enrichment (EE; 91.5×64×159 cm; see **Figure 1B**), which was a large multi-level cage with ramps and access to toys, tubes, chew bones, and Nestlets^®^ (Ancare, Bellmore, NY), or standard laboratory cages (SD; 27×48×20 cm; see **Figure 1C**). Enrichment toys (e.g., small plastic balls, climbing ropes and ladders, swings, bell rollers, chew toys, hammocks, additional tubes/tunnels, Lixit Space Pods, cups, and other small animal hideaways) were switched out twice weekly to maintain novelty in the EE condition.

**Figure 1.**
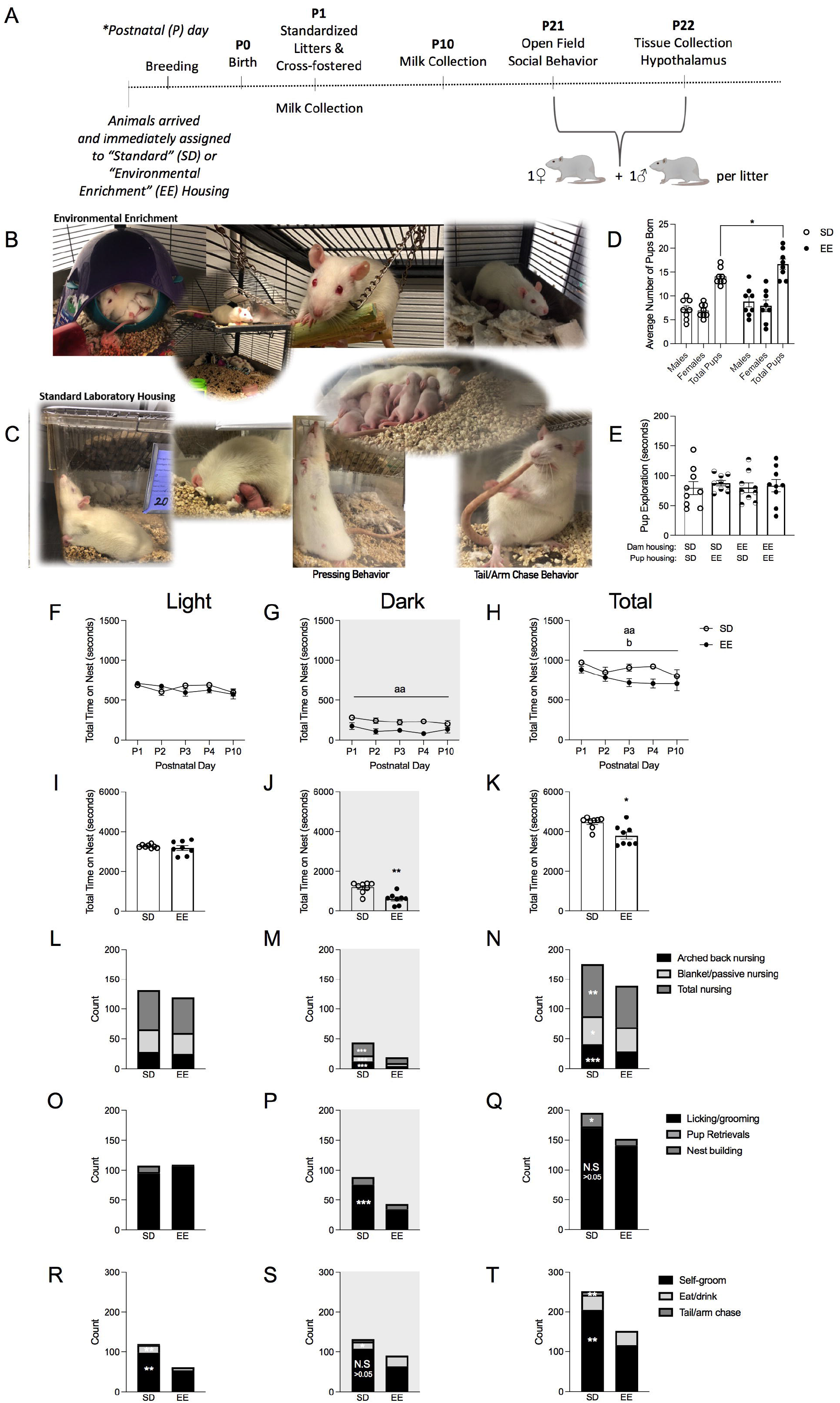
Maternal care behaviors are different between environmentally enriched (EE) and standard housed (SD) Sprague-Dawley rat dams. (A) Timeline of experimental procedures. (B) Representative photographs of EE housing and litters. (C) Representative photographs of SD housing and litters. (D) Average number of pups born (male, female, and total pups) per SD and EE housing group (n = 8). (E) Total time (seconds) SD and EE housed dams spent exploring postnatal day (P)7 alien pups from different housing conditions (n = 9). Total time (seconds) that dams spent on the nest across P1-P4 and P10 in the (F) light, (G) dark, and (H) light + dark periods combined. Total time (seconds) that dams spent on the nest collapsed across P1-P4 and P10 in the (I) light, (J) dark, and (K) light + dark periods combined. Stacked bars depict the frequency of pup directed nursing behaviors (arched back nursing, blanked/passive nursing, total nursing) collapsed across P1-P4 and P10 in the (L) light, (M) dark, and (N) light + dark periods. Stacked bars depict the frequency of other types of pup directed behaviors (licking/grooming, pup retrievals, nest building behaviors) collapsed across P1-P4 and P10 in the (O) light, (P) dark, and (Q) light + dark periods. Stacked bars depict the frequency of maternal self-directed behaviors (self-grooming, eating/drinking, tail/arm chases) collapsed across P1-P4 and P10 in the (R) light, (S) dark, and (T) light + dark periods (n = 8). Data are expressed as mean ± SEM; SD: open circles versus EE: closed circles. *p < 0.05, ** p < 0.01, ***p <0.001, SD versus EE; ^aa^p < 0.01, main effect of housing; ^b^p < 0.05, main effect of postnatal day.

Male rats were paired in SD conditions unless they were breeding. During breeding, they were housed with two females in either EE housing or in larger SD one-level cages (51×41×22 cm) with access to a tube, one chew bone and Nestlets^®^ (Ancare, Bellmore, NY). Approximately two days prior to parturition, dams in the SD condition were individually housed (27×48×20 cm; **Figure 1C**), while a physical divider separated the EE dams within their cage (allowing for auditory, tactile, olfactory, and some visual contact; important components of EE). This separation prevented the mixing of litters. Day of birth was designated as postnatal day (P) 0 and litters were standardized to 10 pups per litter on P1. To dissociate the effects of the pre- and postnatal housing environments, male and female pups from each litter were cross-fostered at this time. To track the housing of origin, pups were marked on their left or right ear to indicate prenatal SD or EE respectively, resulting in the following study group designations: SD-SD, SD-EE, EE-SD, EE-EE. Offspring were maintained in these respective housing conditions until the end of the study on P22. All animal procedures were performed in accordance with the [Author University Blinded] animal care committee’s regulations.

### Milk sample collection

To encourage the accumulation of maternal milk for sample collection, on the mornings of P1 (equivalent to a transitional milk phase) and P10 (mature milk phase), dams were removed from their litter and placed into a clean cage in a separate procedure room for one hour. Pups remained in their regular holding room and were placed into a smaller clean cage positioned on top of a heating pad, to maintain their body temperature. Litters were weighed immediately prior to being returned to their dams and again 2 and 24 hours later, alongside the inspection of milk bands, to monitor their health.

The milk procedure was adapted from a published procedure (Paul et al., 2015). Immediately following the separation period, dams were lightly anesthetized with isoflurane in O2, followed by the administration of 0.2 mL of oxytocin (20 USP/mL i.p). Distilled water was used to moisten the teats and milk obtained by gently squeezing its base to manually expel the sample for collection. A microhematocrit tube was used to collect ~20 ul of sample. The tube was then sealed and placed into a hematocrit spinner and spun for 120 sec at 13, 700 g. Measurements of the separation of the milk into cream and clear layers were taken to calculate percent (%) creamatocrit using the procedures outlined in Paul et al. (2015). The remaining milk collected (~500 ul per animal) was transferred to small centrifuge tubes and stored at −80 degrees Celsius until processing. Collection time took about 10-20 minutes per animal, and dams were returned to their litter as soon as they awoke from anesthesia. This was appropriate as breastfeeding can resume immediately after isoflurane anesthesia since the pharmacokinetics of the compound indicate it is poorly absorbed by infants (Drugs and Lactation Database, 2020; Lee & Rubin, 1993). With respect to oxytocin, its typical half-life (1-6 minutes) is reduced even further during lactation and this drug is also unlikely to affect offspring (Par Pharmaceutical Inc, 2020).

### Milk sample immunoassays

Milk samples (n = 8) were placed onto a Mini Tube Rotator (Fisher Scientific Cat. #88861051) overnight at 4°C to homogenize prior to analysis. Following the standard manufacturer’s instructions, commercially available ELISA kits were used to measure lactose content (Sigma-Aldrich Cat. #MAK017; diluted 1:500; as outlined by Chen et al., 2017; DeRosa et al., 2022), triglycerides (Abcam, Cat. #ab65336; 1:1000 dilution), protein (Pierce™ BCA Protein Assay Kit; Cat. #23227; 1:50 dilution), and immunoglobulin (Ig) A (Bethyl Laboratories, Cat. # E111-102; diluted to 1:1000) levels in milk. To measure corticosterone, the small sample assay protocol (#ADI-900–097, Enzo Life Sciences, Farmingdale, NY) was followed, as recommended by the manufacturer, using a 1:40 dilution (DeRosa et al., 2022). We opted to only evaluate P10 milk samples on these measures because litters were appropriately standardized in size for this time point.

### Microbiome sequencing of milk samples

Another subset of P10 milk samples underwent microbiome community analysis (n = 6). Milk DNA was first extracted using the ZymoBIOMICS ^®^-96 MagBead DNA Kit (Zymo Research, Irvine, CA). The *Quick-16S*™ NGS Library Prep Kit (Zymo Research, Irvine, CA) was used for bacterial 16S ribosomal RNA targeted sequencing and custom 16S primers were utilized to amplify the V3-V4 region (Zymo Research, Irvine, CA). Real-time PCR was then used to prepare the sequencing library and final qPCR fluorescence readings were pooled together according to equal molarity and the final pooled library was cleaned using the Select-a-Size DNA Clean & Concentrator™ (Zymo Research, Irvine, CA), and quantified with TapeStation^®^ (Agilent Technologies, Santa Clara, CA) and Qubit^®^ (Thermo Fisher Scientific, Waltham, WA). Illumina^®^ MiSeq™ with a v3 reagent kit (600 cycles) was used along with a 10% PhiX spike-in to sequence the final library. Unique amplicon sequences, as well as possible sequencing errors and chimeric sequences, were inferred from raw reads using the DADA2 pipeline (Callahan et al., 2016). Uclust from Qiime (v.1.9.1) was used to determine taxonomy assignment and referenced with the Zymo Research Database (Zymo Research, Irvine, CA).

### RNA-sequencing of milk samples

Total RNA (n=4-5 per housing group) was isolated from a subset of P10 milk samples using a previously published procedure (Chen et al., 2017; DeRosa et al., 2022). Milk was centrifuged for 5 minutes at 1,000 g (4°C) and the fat layer isolated. Milk fat was then mixed with an equal volume of PBS by centrifuging at 3,000 rpm (5 min in 4°C), cleaning the fat of debris. The miRNeasy Mini Kit (QIAGEN, Cat. #217004) was used to isolate total RNA following the manufacturer’s directions. A NanoDrop 2000 spectrophotometer (ThermoFisher Scientific) was used to quantify isolated RNA samples, which were then stored at −80°C. For RNA-sequencing (RNA-seq), the quality of the RNA was determined using Bioanalyzer (Agilent technologies, Santa Clara, USA). The cDNA library was compiled using the TruSeq Stranded mRNA kit (Illumina, San Diego, USA), and processed through Tapestation to determine fragment size and DNA concentration. The library was then sequenced on an Illumina NovaSeq 6000 to obtain single-end 100-bp reads. Samples were read at a sequencing depth of approximately 50 million reads. These reads were then aligned to the rn6 genome using the BSgenome.Rnorvegicus.UCSC.rn6 R/Bioconductor package (version 1.4.1). Differentially expressed genes, using a p < 0.05, Benjamini–Hochberg false discovery rate (FDR) corrected, and fold change (FC) > 1.3, were identified using DESeq2 package (Love et al., 2016). Heatmaps were generated using the z-score of rlog-normalized counts and were plotted with the MultiExperiment Viewer (National Library of Medicine, USA). Gene ontology from genes with p < 0.05 or FDR adjusted p <0.05 with a FC > 1.3 was generated using the Database for Annotation, Visualization and Integrated Discovery functional annotation cluster tool (https://david.ncifcrf.gov/). RNA-seq data have been deposited to GEO (GSE200249).

### Maternal care

Maternal behavior observations took place between P1-P4 and again on P10 following the milking procedures (n = 8). Sessions occurred three times daily (7:30, 15:00, 20:00), consisting of six observations resulting in a composite score for each dam and observation period. Dams were evaluated for 1-minute intervals per observation, with at least 5 minutes of no observations occurring between each of the 1-minute bins. Maternal care observations recorded included the frequency of pup-directed behaviors (i.e., dam licking/grooming pups, active/high crouch nursing, passive/low crouch nursing, pup retrieval), self-directed behaviors (i.e., dam eating/drinking, dam self-grooming, dam chasing her arm/tail), and nest building/digging behavior. Total time the dam spent on her nest (seconds) was also recorded (Strzelewicz et al., 2021; Strzelewicz et al., 2019; Connors et al., 2015).

### Open field and social preference tests

On P21, one male and one female offspring per litter from each pre-versus postnatal housing condition was habituated to an open field arena for three-minutes (40 cm × 40 cm × 28 cm; Duque-Wilckens et al., 2020; Williams et al., 2020, n = 7-8). Behavior was recorded and videos scored using an automated behavioral monitoring software program (Cleversys TopScan, Reston, VA) to determine total distance traveled (cm) and percent of time spent in the center of the arena. All equipment was thoroughly cleaned with Quatriside TB between each animal and test. Immediately after the open field habituation period, animals were evaluated in a five-minute social preference test. Using the manual behavioral monitoring program ODLog™ 2.0 (http://www.macropodsoftware.com/), animals were evaluated on their choice to visit a novel inanimate object or a novel same sex and age conspecific, each enclosed within a small wire cup on opposite ends of the arena (Crawley, 2007). Placement of novel rats and objects were interchanged between trials and experimental groups counterbalanced between tests. Animals were recorded as actively investigating when their nose was directed within 2 cm of a containment cup, or it was touching the cup. Percent time in contact with either the novel rat or object was calculated by the formula ([total time with target cup (rat or object) / 300 seconds] *100). Latency (seconds) to approach the novel rat was also recorded (Strzelewicz et al., 2019).

In a separate group of rat dams (n =9), a pup preference test was run using a similar protocol. The purpose of this test was to determine if dams had a social preference for either SD or EE housed P7 pups, which may have impacted maternal care and later offspring social behavior (Champagne et al., 2009; Cromwell, 2011). Dams were assessed on the amount of time they spent exploring alien SD versus alien EE housed pups. One male and one female pup from each housing condition were placed into wire cups situated on opposite sides of an open field arena. The total duration (seconds) that SD and EE dams spent with each housing group across a ten-minute period was reported.

### Western blotting

On the morning of P22, a mixture of ketamine/xylazine (150 mg/kg, i.p./50 mg/kg, i.p.) was used to anesthetize dams and their litters. Maternal blood was collected in EDTA coated tubes following a cardiac puncture and spun at 1,000g for 10 minutes to obtain plasma. Levels of prolactin were determined in undiluted plasma samples using an ELISA (Abcam, Cat.# ab272780). Whole hypothalamus was dissected from offspring and dams, frozen on dry ice, and stored at −80°C for future processing. Hypothalamic tissue was later homogenized and the amount of protein in each sample was determined using the BCA assay (Pierce™ BCA Protein Assay Kit; Cat. #23227). Protein was mixed with 2x Laemmli sample buffer (Bio Rad Laboratories Cat. #1610737) and denatured at 100 °C for 5 minutes. 20μg of protein was loaded into each well of Mini-Protean^®^ gels (Bio Rad Laboratories, Cat. #4568101) and transferred onto nitrocellulose membranes (Bio Rad Laboratories, Cat. #1620147). Membranes were then incubated in a 5% nonfat milk with TBS + 0.05% Tween 20 (TBST) blocking solution for 1 hour at room temperature. Following this, they were washed with TBST and incubated in a 1:1000 dilution of CB1 receptor antibody (Abcam Cat. #ab259323) in TBS solution overnight at 4 °C. The next morning, membranes were washed and incubated in an HRP-conjugated secondary antibody (1:1000, Abcam, Cat., #ab131366), made in 1% nonfat milk with TBS, for 1 hour at room temperature. Membranes were then washed and immersed in a chemiluminescent substrate (Thermo Fisher Scientific, Cat. #34580) for 5 minutes prior to being scanned with a LI-COR C-DiGit Scanner (Model # 3600). After imaging, membranes were stripped (Thermo Fisher, Cat. # 21062) for 15 minutes at 37°C, followed by blocking in 5% nonfat milk with TBST for 1 hour at room temperature. After washing, the membranes were incubated in β-actin (1:1000, Thermo Fisher Scientific, Cat. #MA515739) for 1 hour and imaged again. Densitometry was used to obtain a ratio of CB1/ β-actin in order to quantify protein differences between housing groups.

### Statistical analyses

Statistics were performed using the software package Statistical Software for the Social Sciences (SPSS) version 26.0 (IBM, Armonk, NY) or GraphPad Prism (version 9.0). The dataset was not powered to evaluate sex-differences so male and female animals were evaluated separately (Ordoñes Sanchez et al., 2021). Two-way repeated measure ANOVAs (Housing x Time) were used to compare P1 and P10 milk levels of creamatocrit, which is linearly related to the fat concentration and energy content of milk (Lucas et al., 1978; Paul et al., 2015). This statistical analysis was also used to compare the total time the dam spent on the nest (P1-P4, P10) across the light and dark phases of the circadian cycle.

A paired samples t-test was used to evaluate maternal preferences for P7 SD and EE housed pups. One-way ANOVAs were used to evaluate other measures of maternal care and milk composition (e.g., ELISA data) as a function of housing condition. In rare cases of violations to the assumption of normality (Shapiro-Wilk test), Kruskal-Wallis tests were employed (expressed as *X^2^*). Offspring behavior was assessed using 2 x 2 (prenatal treatment x postnatal treatment) ANOVAs and LSD post hocs were applied except where there were fewer than three levels, in which case pairwise t-tests and Levene’s (applied in the occurrence of unequal variances) were utilized alongside Bonferroni alpha adjustments.

Data are graphically expressed as mean ± SEM. The partial eta-squared (*n_p_*^2^) is also reported as an index of effect size for the ANOVAs (the range of values being 0.02 = small effect, 0.13 = moderate effect, 0.26 = large effect; Miles and Shevlin, 2001).

For the microbiome analyses, composition visualization, alpha-diversity, and betadiversity analyses were performed with Qiime (v. 1.9.1) and statistical comparisons were performed using Kruskal-Wallis tests (Caporaso et al., 2010). To determine taxa that were significantly different between groups, linear discriminant analysis effect size (LEfSe; http://huttenhower.sph.harvard.edu/lefse/) was employed as previously described (Segata et al., 2011; Schellekens et al., 2021). In short, LEfSe creates a model that identifies taxa that are most likely to explain differences between groups through the use of a series of nonparametric tests (Segata et al., 2011).

DESeq2 was used to determine differentially expressed genes based on p< 0.05 or FDR adjusted p<0.05, with a fold change threshold of 1.3 for RNA-seq.

## Results

### Maternal housing condition affects maternal care

One-Way ANOVA showed that EE dams gave birth to larger litters than their SD housed counterparts (SD: 14.0±0.60 vs. EE: 16.63±1.05; F(1, 14) = 4.173, p = 0.048, *n_p_*^2^ = 0.252; **Figure 1D**); all litters were standardized to 10 pups on P1. On P7, a social preference test showed that EE and SD dams investigated all pups to the same extent, regardless of the pups housing origin (t(17) = 0.808, p = 0.430; **Figure 1E**).

A repeated measures ANOVA suggested that the time dams spent on the nest did not change as a function of postnatal day across the light and dark periods (p >0.05; **Figure 1F, G**). However, when these two observation periods were collapsed together a significant main effect of postnatal day emerged (F(4, 56) = 2.740, p = 0.037, *n_p_*^2^ = 0.164; **Figure 1H**) with more time being spent on the nest on P3 (p = 0.013) and P4 (p < 0.001), an effect driven by the SD group. Indeed, both the dark, and the total light + dark periods combined revealed a significant main effect of housing condition in that SD dams spent more time on the maternal nest than EE dams (dark: F(1, 14) = 16.987, p = 0.001, *n_p_*^2^ = 0.548; **Figure 1G**; total light + dark: F(1, 14) = 9.839, p = 0.007, *n_p_*^2^ = 0.413; **Figure 1H**). This general pattern persisted when the total time dams spent on the nest was summed into a composite score across the postnatal observation days. Again, SD dams were shown to spend more time on the nest than EE mothers (light: p>0.05, **Figure 1I**; dark: *X^2^*(1) = 7.456, p = 0.006, **Figure 1J**; total light + dark: *X^2^*(1) = 4.864, p = 0.027; **Figure 1K**). During our maternal care observations, we did not plan to quantify “press posture” (Gaskill & Pritchett-Corning, 2015; Cramer et al., 1990; Ratuski & Weary 2021) but we subjectively noted its presence in SD dams when it was noticed during our study. A representative photo of this posture can be found in **Figure 1C**. We did not observe this behavior in any EE dams.

Maternal nursing postures did not differ as a function of housing in the light phase (p>0.05; **Figure 1L**). However, the number of total nursing postures observed were significantly lower in EE dams during the dark phase (high arched back: F(1, 14) = 24.462, p = 0.001, *n_p_*^2^ = 0.636; passive/blanket: *X*^2^(1) = 4.498, p = 0.034; total nursing: F(1, 14) = 17.996, p = 0.001, *n_p_*^2^ = 0.562; **Figure 1M**) and in the total combined light + dark phases (high arched back: F(1, 14) = 10.772, p = 0.005, *n_p_*^2^ = 0.435; passive/blanket: F(1, 14) = 5.296, p = 0.037, *n_p_*^2^ = 0.274; total nursing: *X*^2^(1) = 9.289, p =0.002; **Figure 1N**).

Maternal nest building bouts were significantly increased in SD dams during the light phase (*X^2^*(1) = 7.323, p = 0.007; **Figure 1O**). SD dams also licked/groomed their pups more frequently in the dark phase (F(1, 14) = 22.257, p = 0.001, *n_p_*^2^ = .614; **Figure 1P**). However, the total amount of licking and grooming that EE and SD pups received across the total light + dark phases did not differ (p>0.05; **Figure 1Q**), while increased nest building was sustained in SD dams (*X*^2^(1) = 5.370, p = 0.020; **Figure 1Q**).

During the light phase, maternal self-directed grooming and eating/drinking behaviors were elevated in SD dams (maternal self-grooming: F(1, 14) = 6.077, p = 0.027, *n_p_*^2^ = 0.303; eating/drinking: F(1, 14) = 4.833, p = 0.045, *n_p_*^2^ = 0.257; **Figure 1R**). SD dams also displayed a higher number of repetitive tail/arm chase behaviors (see **Figure 1C** for photograph) across the nychthemeron (light: *X*^2^(1) = 6.536, p = 0.011; dark: *X*^2^(1) = 6.303, p = 0.012, **Figure 1S**; total light + dark: *X*^2^(1) = 6.792, p = 0.009; **Figure 1T**) and higher self-grooming levels when the light + dark periods were combined (F(1, 14) = 9.914, p = 0.007, *n_p_*^2^ = 0.415; **Figure 1T**).

### Environmental enrichment increases lactation quality in terms of protein expression of prolactin, creamatocrit, and triglycerides

Given that SD dams spent significantly more time on the nest feeding their litters than EE dams, and that EE dams had higher plasma concentrations of the breastmilk-producing hormone prolactin (F(1, 13)= 5.69, p=.034, *n_p_*^2^= .322; **Extended Data Figure 1-1**), we evaluated whether there were differences in milk quality between the two housing conditions (see **Figure 2A** for photograph of milk collection procedure). There was a main effect of postnatal day for % creamatocrit, which is directly proportional to the fat concentration and energy content of milk (Lucas et al., 1978; Paul et al., 2015). These proportionally related measures decreased in both housing groups between P1 and P10 (F(1, 14) = 23.607, p = 0.001, *n_p_*^2^ = 0.001; **Figure 2B, C, D**) The ratio of different milk contents changes over the course of lactation in rats, and fat in particular decreases with time (Keen et al., 1981). There were no significant housing effects in the concentration of protein, lactose, corticosterone, or IgA in P10 milk (p >0.05; **Figure 2E, F, G, H**). However, triglyceride levels were significantly higher in the milk of EE compared to SD housed dams (F(1, 14) = 9.314, p = 0.009, *n_p_*^2^= 0.400; **Figure 2I**).

**Figure 2.**
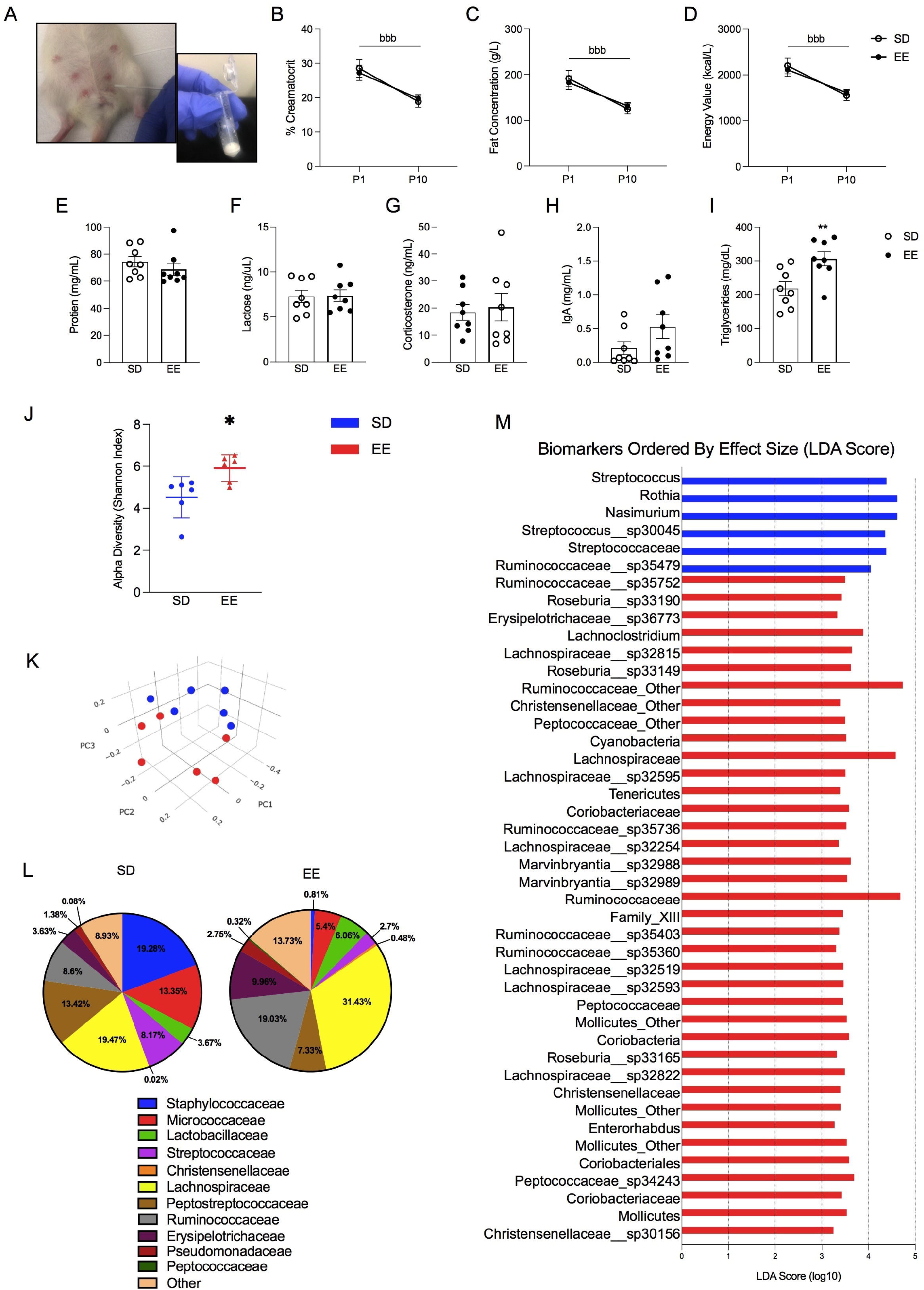
Nutritional profile of milk and microbiome community distribution in environmentally enriched (EE) and standard housed (SD) Sprague-Dawley rat dams. (A) Photograph depictions of maternal milk collection. Maternal milk concentrations (n = 8) of (B) % creamatocrit, (C) fat (g/L), (D) energy value (kcal/L), (E) protein (mg/mL), (F) lactose (ng/μL), (G) corticosterone (ng/mL), (H) IgA (mg/mL), and (I) triglycerides (mg/dL). Microbiome sequencing (n = 6) is demonstrated with (J) Alpha diversity along the Shannon Index and (K) Beta diversity using principle coordinate analysis (PCoA). This plot was created using the matrix of pair-wise distance between samples determined by Bray-Curtis dissimilarity using unique amplicon sequencing variants. Each dot represents an individual microbial profile. Samples that are closer together are more similar, while samples that are dissimilar are plotted further away from one another. (L) Microbial composition of taxonomy in maternal milk at the family level for SD and EE-housed dams. (M) Microbiome biomarkers plot. Taxa identified as significantly more abundant in the milk of the housing group where a bar appears; SD mothers (blue bars) and EE mothers (red bars). Significance was determined by LEfSe analysis, which identified taxa with distributions that were statistically significant (p < 0.05) and where the effect size (LDA score) was greater than 2. Data are expressed as mean ± SEM; SD: open circles versus EE: closed circles. **p <0.01, SD versus EE; ^b^p < 0.05, main effect of postnatal day.

### Environmental enrichment increases the microbiome diversity of maternal milk which has a higher abundance of bacterial families related to body weight and energy metabolism

Housing condition also contributed to significant differences in the composition of the milk microbiome (**Figure 2J-M)**. EE milk contained greater species diversity compared to SD milk, as indicated by alpha diversity along the Shannon index ((*X*^2^(1) = 5.77, p = 0.016; **Figure 2J)** and Bray-Curtis dissimilarity measurement of beta diversity (R = 0.2944, p = 0.02; **Figure 2K**). Please see **Extended Data Figures 2-1 and 3-1** for the Cladogram of milk biomarkers and taxonomy heatmap respectively. Overall, LEfSe analysis revealed 44 discriminative taxa between our housing groups, 38 of which were more highly expressed in the milk of EE dams. Specifically, milk from EE dams demonstrated a significantly greater abundance of the phylum *Tenericutes* (LDA effect size = 3.39, p = 0.03; **Figure 2J**). Additionally, *Christensenellaceae* (LDA effect size = 3.39; p = 0.007), *Peptococcaceae* (LDA effect size = 3.499; p = 0.02), *Coriobacteriaceae* (LDA effect size = 3.59; p=0.006), *Lachnospiraceae* (LDA effect size = 3.32; p = 0.007), *Ruminococcaceae* (LDA effect size = 4.73; p = 0.02) and *Erysipelotrichaceae* (LDA effect size = 3.33; p = 0.02) were higher at the family level (**Figure 2L, M**) in EE mothers. Milk from SD dams had greater levels of *Streptococcaceae* (LDA effect size = 4.38; p = 0.02; **Figure 2L, M).**

### RNA-sequencing identified several maternal milk transcriptomic pathways that were differently affected by housing condition

Based on our identified immunoassay targets and microbiome evaluations, we broadened our analyses to the whole genome context by evaluating transcriptomic profiles of SD and EE milk samples which were characterized using RNA-seq. A total of 756 genes were differently expressed (p<0.05, FC>1.3) between housing conditions with 110 genes meeting the significant threshold padj<0.05 (**Figure 3A, Table 3-1**). First, we targeted the ontology analysis towards genes that contribute to maternal milk nutrition (Strucken et al., 2015). RNA-seq revealed several differently expressed genes involved with milk triglyceride and nutrient transport downregulated by EE, including *Ghr* (p = .024, FC = −0.671), *Igf1* (p = 0.03, FC = −0.671), *Slc27a4* (p= 0.032, FC = −0.447), *Gpat4* (p<.001, FC = −0.613), and *Cs1s2b* (p = 0.009, FC = −0.838; Velmala et al., 1995; **Figure 3B, C**). We broadened the scope of our analyses further to include gene pathways related to oxytocin (Quintana et al., 2019), glucocorticoid receptor (GR) signaling (e.g., events that occur when glucocorticoids bind to the GR receptor; (Oakley & Cidlowski, 2013), GR binding (e.g., genes which bind the glucocorticoid receptor; Polman et al., 2012), and epigenetic modifiers (Zhu et al., 2016; **Figure 3D, E, F, G**) given their crucial implication in milk production and offspring development (Ozkan et al., 2020; Zhang et al., 2013). Heatmap clustering showed that EE rats displayed a major downregulation of genes encoding for triglycerides and nutrient transport **(Figure 3B, C)**, while genes related to oxytocin and GR **(Figure 3D, E)** signaling were mostly upregulated compared to SD rats. Another cluster of genes related to glutamate/GABA signaling (Gray et al., 2018) were mostly downregulated by EE (**Figure 4-1A**). GR binding genes, epigenetic modifiers, and prolactin-signaling-related genes (Radhakrishnan et al., 2012) were instead equally regulated in both directions (**Figure 4-1B**).

**Figure 3.**
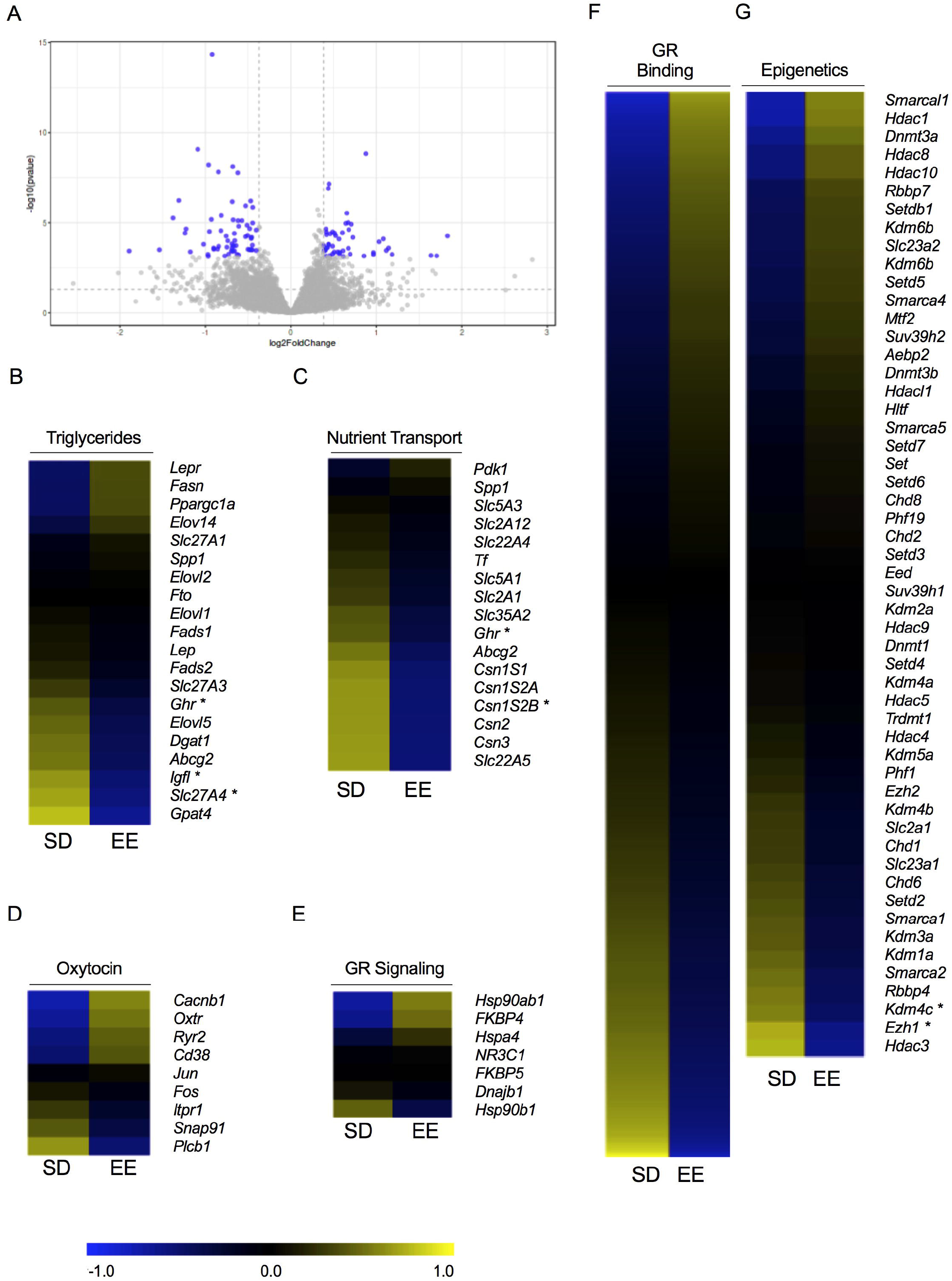
Transcriptomic analyses of P10 milk samples from rat mothers living in environmental enrichment (EE) or standard housing (SD). (A) Volcano plot depicting the distribution of 756 genes based on log2 fold change and -log10 p values. Each grey dot is a gene, and 110 dots highlighted in blue represented genes that displayed the highest magnitude of significance (padj<0.05, FC>1.3). Heatmaps of differentially expressed genes related specifically to milk (B) triglyceride concentration, (C), nutrient transport, (D), oxytocin signaling, (E) GR signaling, (F) GR binding, and (G) epigenetics. Gene expression is represented with the log2 transformation of counts recorded with a z-score based on the average across experimental groups. Data are expressed as *p <0.05 or **adjusted p <0.05, with FC>1.3. GR-glucocorticoid receptor.

### Enriched offspring had heavier body weights and increased social behavior which was lost by cross-fostering into standard housing in a sex-specific manner

A two-way (prenatal x postnatal) ANOVA revealed a main effect of postnatal experience for male and female offspring P21 body weights (males: *X*^2^(1) = 9.562, p = 0.002; females: *X*^2^(1) = 11.733, p = 0.001, **Figure 4A, B, C**). Postnatal enrichment housing resulted in significantly higher body weights than SD housing (males SD: 40.43±1.10g versus EE: 50.86±2.01g; females SD: 40.191±0.99g versus EE: 50.50±2.14g).

**Figure 4.**
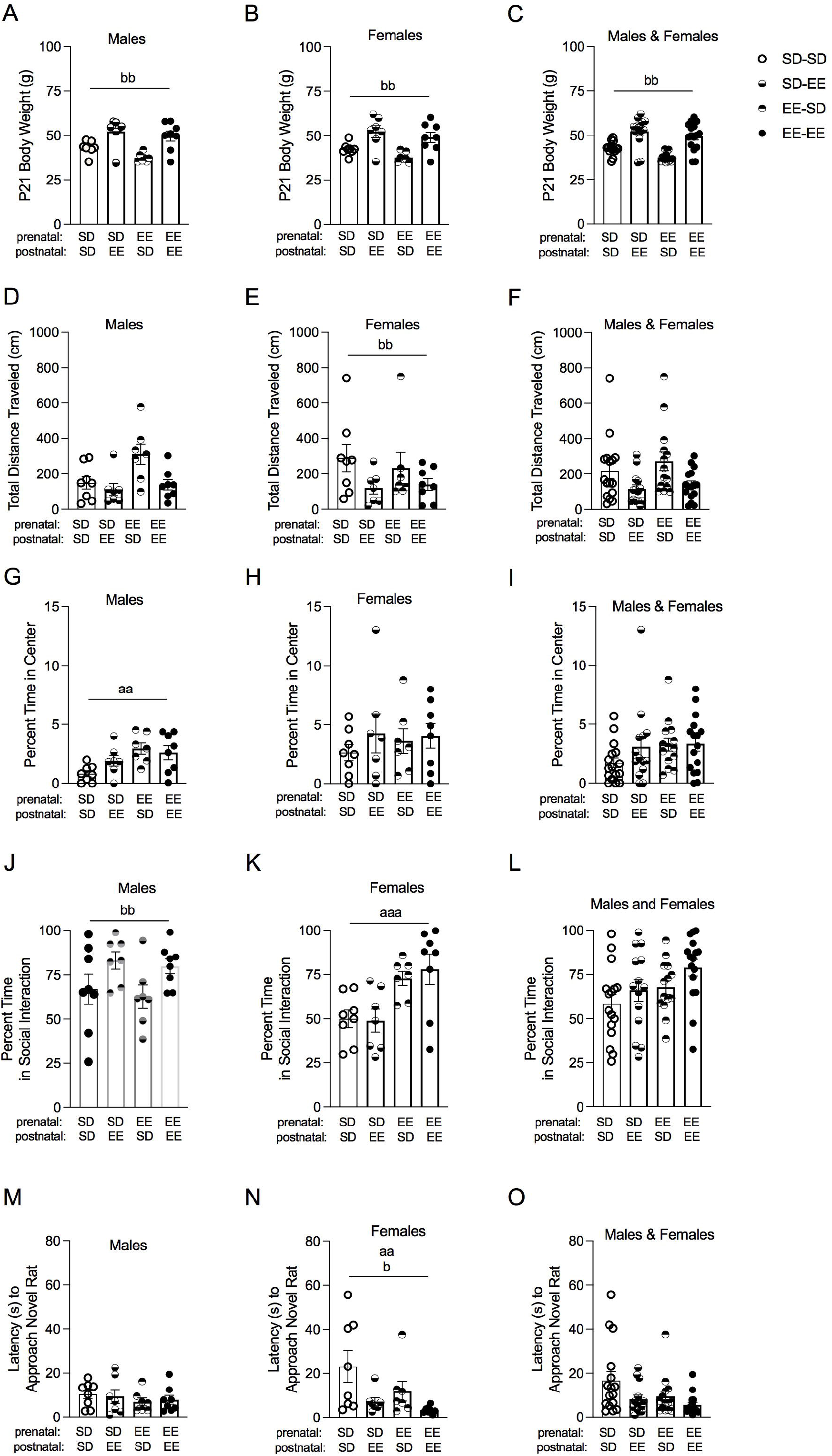
Juvenile offspring physiology and behavior following housing in environmentally enriched (EE) or standard housed (SD) laboratory conditions. Data for (left-side) male, (middle) female, and (right-side) male and female Sprague-Dawley rats combined for (A, B, C) P21 body weights (grams). (D, E, F) Total distance traveled (cm), and (G, H, I) percent of time spent in the center of an open field. (J, K, L) Percent of time spent in social interaction, and (M, N, O) latency (seconds) to approach a novel rat in a social preference test (n = 7-8). Data are expressed as mean ± SEM; SD: open circles versus EE: closed circles. ^aa^p < 0.01, ^aa^p < 0.001, main effect of prenatal experience (SD versus EE); ^b^p < 0.05, ^bb^p < 0.01, main effect of postnatal experience (SD versus EE).

There was a main effect of postnatal experience for total distance traveled in that SD female offspring traveled more than EE females (SD: 262.98 ± 56.69 cm vs EE: 130.41 ± 22.76 cm; *X*^2^(1) = 3.882 p = 0.049; males: p>0.05; **Figure 4D, E, F**). A main effect of prenatal experience was found for male animals (*X*^2^(1) = 7.618, p = 0.006; females: p>0.05; **Figure 4G, H, I**). Male EE (2.76% ± 0.375) offspring spent a significantly higher percentage of time in the center of the arena compared to SD males (1.32% ± 0.285), though the overall times were quite low.

For the percent of time spent in social interaction, there was a main effect of postnatal experience for male offspring in that postnatal enrichment housing increased social interest (SD: 64.88±5.36 vs EE: 81.34±3.09; F(1, 26) = 6.840, p = 0.015, *n_p_*^2^ = 0.208; **Figure 4J, L**); the enrichment effect was blocked for animals cross-fostered into SD housing (**Figure 4J)**. For females, the EE prenatal experience increased social engagement level (SD: 49.44±3.88 vs EE: 75.54±4.91; F(1, 26) = 16.140, p = 0.001, *n_p_*^2^ = 0.383; **Figure 4K, L**). Both prenatal (SD = 15.66 ± 4.37 and EE = 7.30 ± 2.34; *X*^2^(1) = 8.073, p = 0.004) and postnatal (SD = 17.90 ± 4.48 and EE = 5.06 ± 1.02; *X*^2^(1) = 4.047, p = 0.044;) enrichment experience decreased the latency for female offspring to approach the novel rat (**Figure 4M, N, O**).

We assessed the expression of the endocannabinoid receptor CB1 in the hypothalamus of mothers and their offspring given its role in maternal behavior (Schechter et al., 2012) and in the development of social behavior (Argue et al., 2017). While there were no housing-associated differences in CB1 in the offspring (p> 0.05; **Extended Data Figures 5A, B, C**), EE dams exhibited significantly greater expression of CB1 compared to saline dams (*X*^2^(1) = 4.45, p = 0.035; **Extended Data Figure 5D**).

## Discussion

In the present study we demonstrate that EE can change the nutritional and microbial profiles of maternal milk in addition to affecting maternal behavior and offspring development. Our results shed light on the multidimensional impact that EE confers on the pre- and neonatal environment and calls attention to the implications of laboratory housing in developmental animal research. Overall, we observe that cross-fostering blocks the beneficial effects of EE housing on (male) offspring social behavior and (male and female) body weights, implicating maternal milk quality as a driving mechanism (see **Figure 5**). While EE was associated with greater bodyweights and sociability in both sexes, the finding that EE dams spent less time on the nest may seem contradictory. However, laboratory rats will spend time away from the nest when given the opportunity, which facilitates more efficient nursing (Ratuski & Weary, 2021).

**Figure 5.**
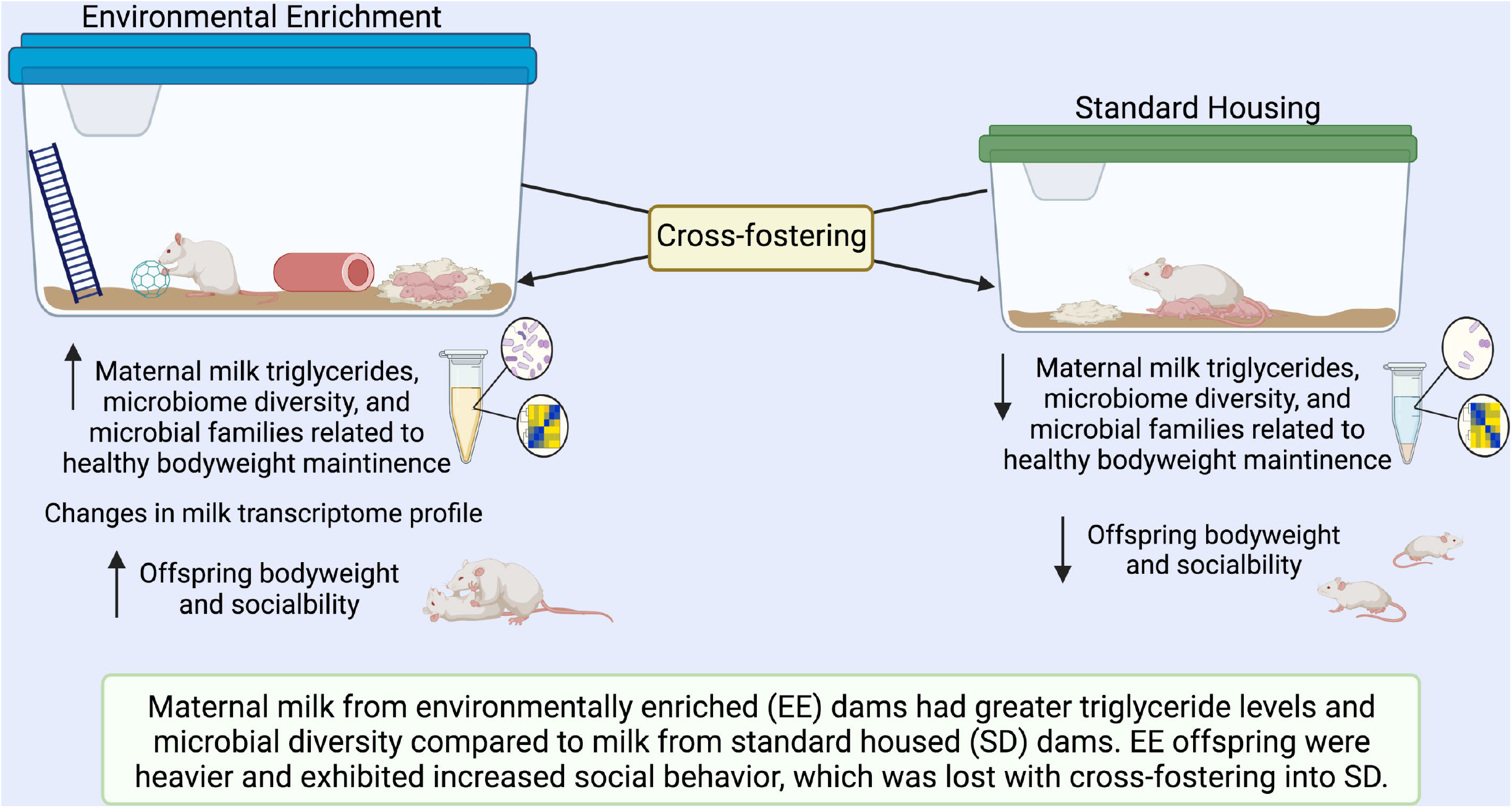
Summary of main results demonstrating that cross-fostering into standard laboratory housing blocks the beneficial effects of environmental enrichment on offspring social behavior and body weight, implicating maternal milk quality as a driving mechanism.

A reduced number of nursing posture displays, coupled with a general increase in time away from offspring, may suggest that EE dams are more efficient at nursing while on the nest. This is further supported by EE-associated increases in circulating prolactin, which promotes milk biosynthesis (Cregan et al., 2001). SD dams may instead need to shift between nursing postures more frequently in order to maintain an active arched back posture, given the extended periods they spend nursing their young. This may artificially increase the amount of nursing postures they display, alongside the timing of maternal behavior observations (e.g., scoring more frequently in the light or dark phases) which can modulate nursing activity level (Peña & Champagne, 2013). Notably, EE and SD dams demonstrated similar amounts of pup licking and grooming despite the differences in time spent on the nest, further supporting the idea that EE dams are more efficient with their care (Welberg et al., 2006; Connors et al., 2015; Strzelewicz et al., 2019; Sale et al., 2004). Higher triglyceride levels in EE milk may compensate for the reduced time on the nest, while SD mothers spend more time nursing to make up for lower milk fat content. Alternatively, their milk quality is reduced because they spend more time feeding as they have limited opportunities to distance from their pups and “re-charge”. This is supported by the greater offspring bodyweights of pups that received postnatal EE.

Greater bodyweights are generally associated with better health in wild animals (Barnett, 1958; Barnett & Dickson, 1984), especially at the age of weaning (Huber et al., 2002; Storey and Snow, 1987). Moreover, heavier adult male wild rats were better integrated socially within their colony (Barnett & Dickson, 1984). These results underscore the translational accuracy of EE used in the present study. The increased bodyweights of EE pups may in part be explained by differences in maternal milk quality. Although we did not find differences in the amount of lactose, protein, or IgA in the milk of EE and SD dams, we did find greater triglyceride levels in EE milk at P10. In support of this finding, Chen et al., (2017) observed that mouse pups fostered to dams with greater levels of milk triglycerides weighed significantly more at P12 than pups that were fostered to control mice. Interestingly, while other studies have also demonstrated the role of increased fat consumption on milk triglyceride levels (Ward et al., 2021; Mohammad et al., 2014) our EE and SD dams did not differ in the composition of their diet or bodyweights. Notably, SD dams ate and drank more than EE mothers suggesting that EE positively contributes to milk triglycerides through a different mechanism.

We further supported our finding regarding increased triglycerides in EE milk with microbiome sequencing. Bacteria in maternal milk prime the infant gastrointestinal tract, which can affect its maturation and the future metabolism of nutrients and bacteria (reviewed in depth in Macpherson et al., 2017). Results of microbiome sequencing demonstrated a robust effect of EE on bacterial diversity in milk samples. For example, milk from EE dams had a significantly higher abundance of *Christensenellaceae, Peptococcaceae, Lachnospiraceae, Ruminococcaceae,* and *Erysipelotrichaceae,* all of which can directly influence lipid metabolism and body mass index through their manipulation of short-chain fatty acids (Bridgewater et al., 2017; Vacca et al., 2020; Greiner & Bäckhed, 2011; Waters & Ley, 2019). Furthermore, milk from EE dams had greater levels of *Coriobacteriaceae,* which contributes to lipid metabolism (Liu et al., 2018), as well as glucose and steroid metabolism in the gut (Clavel et al., 2014). Although increased *Coriobacteriaceae* have been found in the ceca of mice exposed to stress (Bangsgaard Bendtsen et al., 2012), it is unlikely our EE dams were significantly more stressed than our SD dams given the lack of significant differences in milk corticosterone and dam body weights. In addition to finding enhanced microbiome diversity in milk from EE dams, microbiome sequencing revealed significantly greater levels of taxa from the *Streptococcaceae* family in SD dams. Excess expression of this bacterial family in the infant gut has been tied to GI-related issues like dyspepsia and rotavirus infections (Chen et al., 2017; Sohail et al., 2021) which can negatively impact infant growth. Thus, EE may enhance offspring development by altering the microbiome profile of milk that not only promotes the colonization of healthy bacteria, but also discourages exposure to potentially harmful bacteria that can hinder offspring development.

In addition to microbial differences, milk from SD and EE dams demonstrated significant differences in their transcriptomic profiles. Although several of the reported genes related to milk triglyceride and nutrient transport were significantly downregulated in EE dams despite being lipolytic, the sustained activation of genes related to the *Ghr*/*Igf1* feedback loop has previously been shown to reduce maternal milk quality over time in rats (Lékó et al., 2017). Therefore, the reduced expression of these genes in the milk from EE dams may be reflective of more effective endocrinological signaling that allows the mother to return to a basal state. In SD dams however, the constant activation of *Ghr* may be from spending more time on the nest, which compromises milk quality over the course of lactation. Other genes that were differentially regulated between SD and EE milk samples were those related to GR binding, epigenetic modifications, and glutamate/GABA signaling, all of which have considerable implications in offspring brain development and behavior. Together, the results from RNA-seq offer new insight into how housing condition uniquely affects the milk transcriptomics. These data open a door to potential genomic targets and broader networks that are implicated in environmental enrichment models. The future validation of individual genes within these identified pathways through the use of additional techniques like qPCR, will be advantageous in delineating the directionality (i.e., upstream vs downstream) of these networks, and may help reconcile some of the differences observed in offspring physiological and behavioral outcomes.

Our observation of increased sociability in EE offspring is in line with several previous studies (Morley-Fletcher et al., 2003; Peña et al., 2006; Schneider et al., 2006; Connors et al., 2014), although these studies utilized EE to rescue social impairments following early life insults. In saline treated control rats, EE was associated with greater time spent in social interaction (Connors et al., 2014), suggesting that EE is not just protective but can promote sociability. We expand on these findings by revealing sex- and time-specific effects of EE on social preference behavior in healthy offspring. Research assessing the effects of EE on healthy populations is warranted and positively contributes to the translatability of this housing model (Kentner et al., 2019b; Kentner et al., 2021). Cross-fostering pups between housing conditions after parturition revealed that prenatal enrichment increased sociability in females, while postnatal enrichment increased sociability in males. In addition to differences in maternal care, it is plausible that consumption of different milk microbiome profiles may have directly influenced offspring behavior. While the relationship between the gut microbiome and social behavior is well-established (Archie et al., 2015), less is understood about *how* these bacterial taxa exert their effects on brain development and function once colonized in gut. One study demonstrated the necessity of the HPA-axis and neuronal activation in the hypothalamus and hippocampus in rescuing social behavior in germ free adult male mice (Wu et al., 2021). Since germ-free mice are completely devoid of microbiome, this begs the question of whether bacterial taxa can contribute to behavior on a spectrum, and the results of the present study suggest that they may. However, without an assessment of offspring duodenum taxa, we cannot conclude that the gut microbiome profile of our SD and EE offspring differed.

We examined the expression of hypothalamic CB1 in our offspring since this receptor mediates social development in both males and females (Schechter et al., 2012; Argue et al., 2017). Moreover, CB1 in the neonatal brain is associated with the initiation of suckling (Fride et al., 2001). Blocking CB1 function in lactating dams can impair maternal care, prevent pup weight gain, and modulate the social development of offspring (Schechter et al., 2012). Together, these studies suggest that both maternal and offspring CB1 signaling are crucial elements to the lactational period and postnatal offspring development. Although offspring hypothalamic CB1 expression did not differ between our housing groups, EE dams had a significantly greater expression of hypothalamic CB1 compared to saline dams. This supports the involvement of CB1 in the EE-associated changes in maternal nurturance and milk quality, although it is unknown if hypothalamic CB1 activity modulates lactational quality directly. Nonetheless, these findings demonstrate the importance of considering maternal brain physiology in the mediation of offspring developmental outcomes.

While the physiological mechanisms that contribute to time- and sex-specific differences in behavior among EE offspring need further elucidation, the present results shed light on maternal care and maternal milk quality as pathways of interest for future studies to explore. Prenatal measures, such as the hormonal milieu during pregnancy, may also be of interest. EE dams gave birth to larger litters, and this may indicate that the benefits of EE can manifest far before parturition. Presumably, the compounding effects of maternal behavior and milk quality contribute to expedited plasticity in the developing brain (Cancedda et al., 2004; Baroncelli et al., 2010).

## Conclusions

The efficaciousness of the “gold standard” housing cages in animal research has recently been called into question (Kentner et al., 2019b; Olsson & Dahlborn, 2002; Prendergast et al., 2014; Kentner et al., 2021). Results of the present study bolster the argument that EE housing conditions encapsulate a more naturalistic environment than SD, especially with regard to maternal behavior and development (Zhao et al., 2021; Connors et al., 2015; Ratuski & Weary 2021). Recent work has shown running wheel activity to alter milk quality in terms of specific inflammatory molecules such as leukocyte inhibitory factor, CXCL1, and CXCL2; however maternal care was not affected (Taki et al., 2020). This suggests there may be something special in the qualia of the EE condition that extends beyond increased physical activity in terms of its contribution to maternal-neonatal interactions. Overall, we expand on previous studies that have highlighted the beneficial effects of EE on laboratory rodents, by demonstrating the multifaceted impact of EE on maternal behavior, physiology, and offspring social behavior. These results suggest that the maternal environment contributes to notable, long-term changes in offspring development by more efficient maternal behavior and improved milk quality. Rodent models of breastfeeding are advantageous in teasing apart the mechanisms by which this fascinating substance exerts its influence on brain development and behavior.

## Funding Sources

This project was funded by NIMH under Award Number R15MH114035 (to A.C.K).

## Conflict of Interest

The authors report no conflict of interest.

## Acknowledgements

The authors wish to thank Mary Erickson and Sahith Kaki for technical support and Dr. Theresa Casey for advice during the early phases of this study. The authors would also like to thank the MCPHS University School of Pharmacy and School of Arts & Sciences for their continual support, and the University of Massachusetts Boston where HD is a graduate student. Visual abstract made with BioRender.com The content is solely the responsibility of the authors and does not necessarily represent the official views of any of the financial supporters.

## Data Availability

RNA-seq data have been deposited to GEO (GSE200249). DEGs are included in Supplementary Datasets. All other data are available upon request.

## Code Availability

There is no code associated with this work.

---

**Extended Data Figure 1-1. Maternal plasma concentrations of prolactin (ng/mL) for dams housed in environmental enrichment or standard laboratory housing.** Data are expressed as mean ± SEM, *p <0.05, main effect of housing (SD vs EE), n = 7. SD (open circles) or EE (closed circles).

**Extended Data Figure 2-1. Cladogram of milk biomarkers associated with housing condition** (determined by LEfSe). Diameter of the nodes indicates relative abundance of taxa for SD (green) and EE (red) samples. Placement indicates the classification of taxa, where nodes decrease in rank the closer to the center of the diagram.

**Extended Data Figure 3-1. Taxonomy heatmap demonstrating the microbial composition of samples at the species level with the top fifty most abundant species identified.** The colored bar at the top indicates housing condition (blue = SD, red = EE). Each row represents the abundance for each taxon, with the taxonomy ID shown on the right. Each column represents the abundance for each sample.

**Extended Data Figure 4-1. Heatmaps of genes related to glutamate/GABA and prolactin signaling.** Gene expression is represented with the log2 transformation of counts recorded with a z-score based on the average across experimental groups. Data are expressed as *p <0.05 and FC>1.3, or as ***FC>1.3 only.

**Extended Data Figure 5-1. Hypothalamic CB1 and densitometric ratios for offspring and their dams housed in environmental enrichment or standard laboratory housing.** (A) Males, (B) females, (C) males and females collapsed, and (D) mothers housed in SD (open circles) or EE (closed circles) housing conditions. Data are expressed as mean ± SEM, n = 7-8. *p <0.05, main effect of housing (SD vs EE).

**Extended Data Table 3-1. Genes identified with DESeq analysis.** List of genes that significantly differed based on a padj < 0.05 and FC > 1.3 as indicated in Figure 3A.

## Notes

### Competing Interest Statement

The authors have declared no competing interest.

### Summary of Updates

We have evaluated transcriptomic changes in maternal milk quality at the genome-wide level. The associated RNA-sequencing analyses have been added to the manuscript and figures.

