## Supplementary figures and images for "Milking it for all it’s worth: The effects of environmental enrichment on maternal nurturance, lactation quality, and offspring social behavior"

### Extended Data Figure 1-1

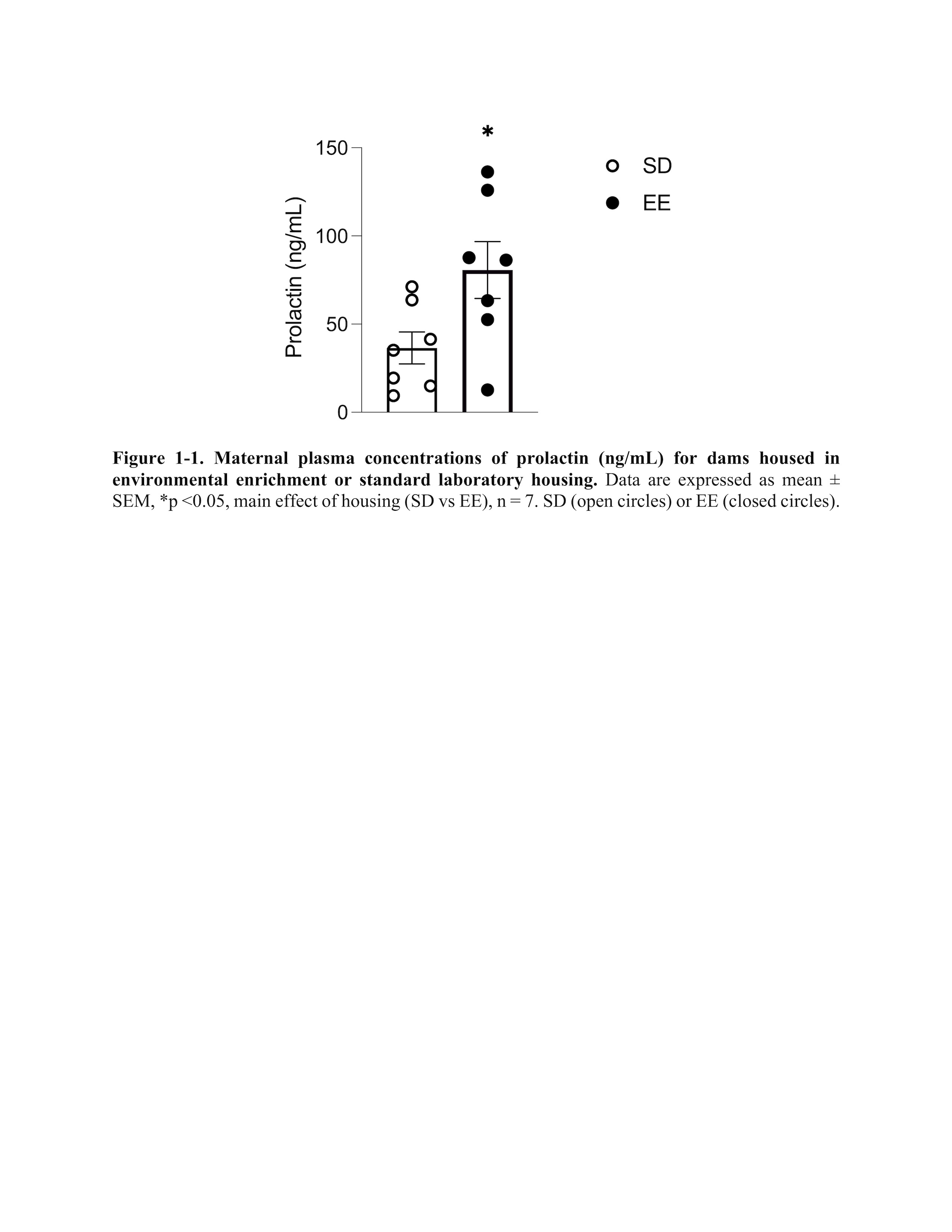

### Extended Data Figure 2-1

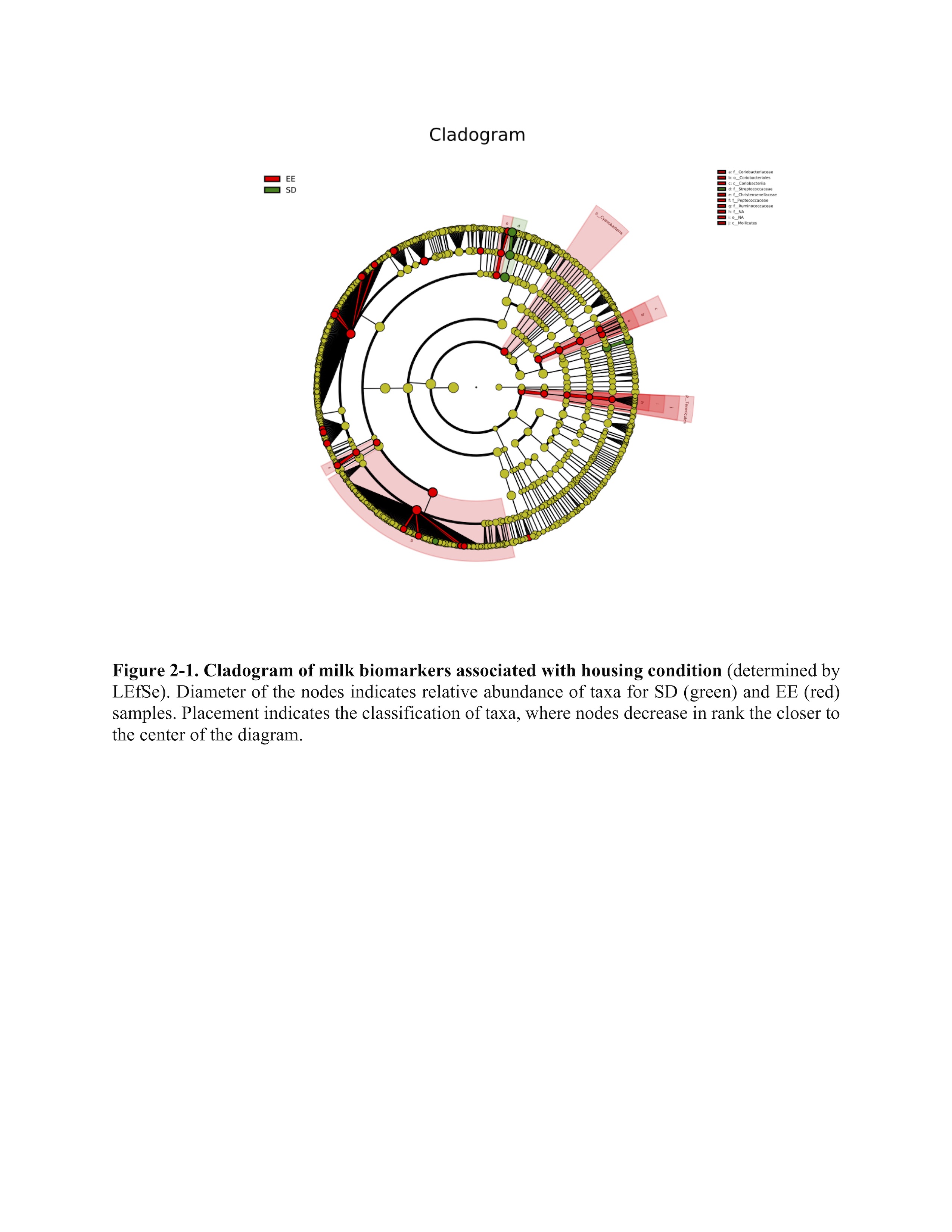

### Extended Data Figure 3-1

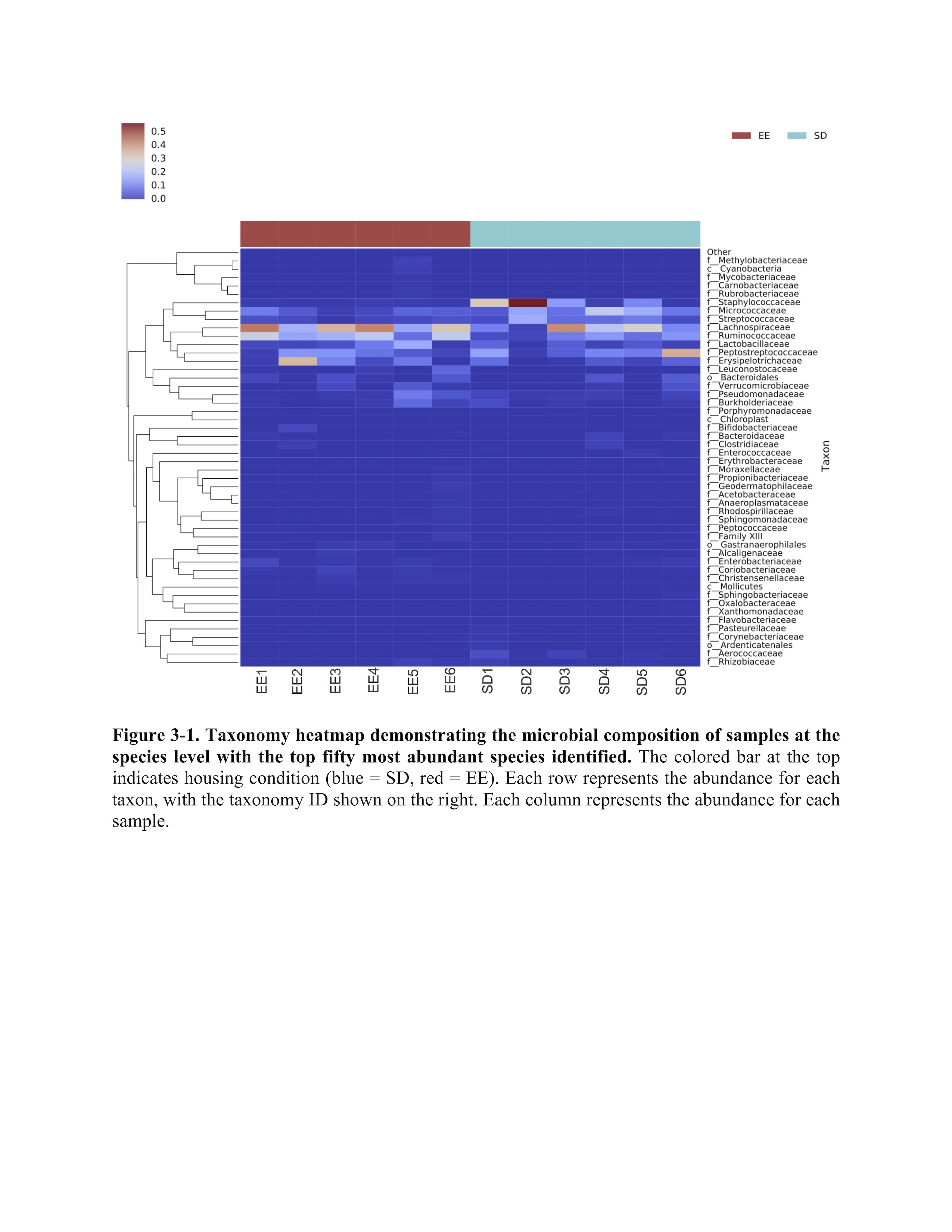

### Extended Data Figure 4-1

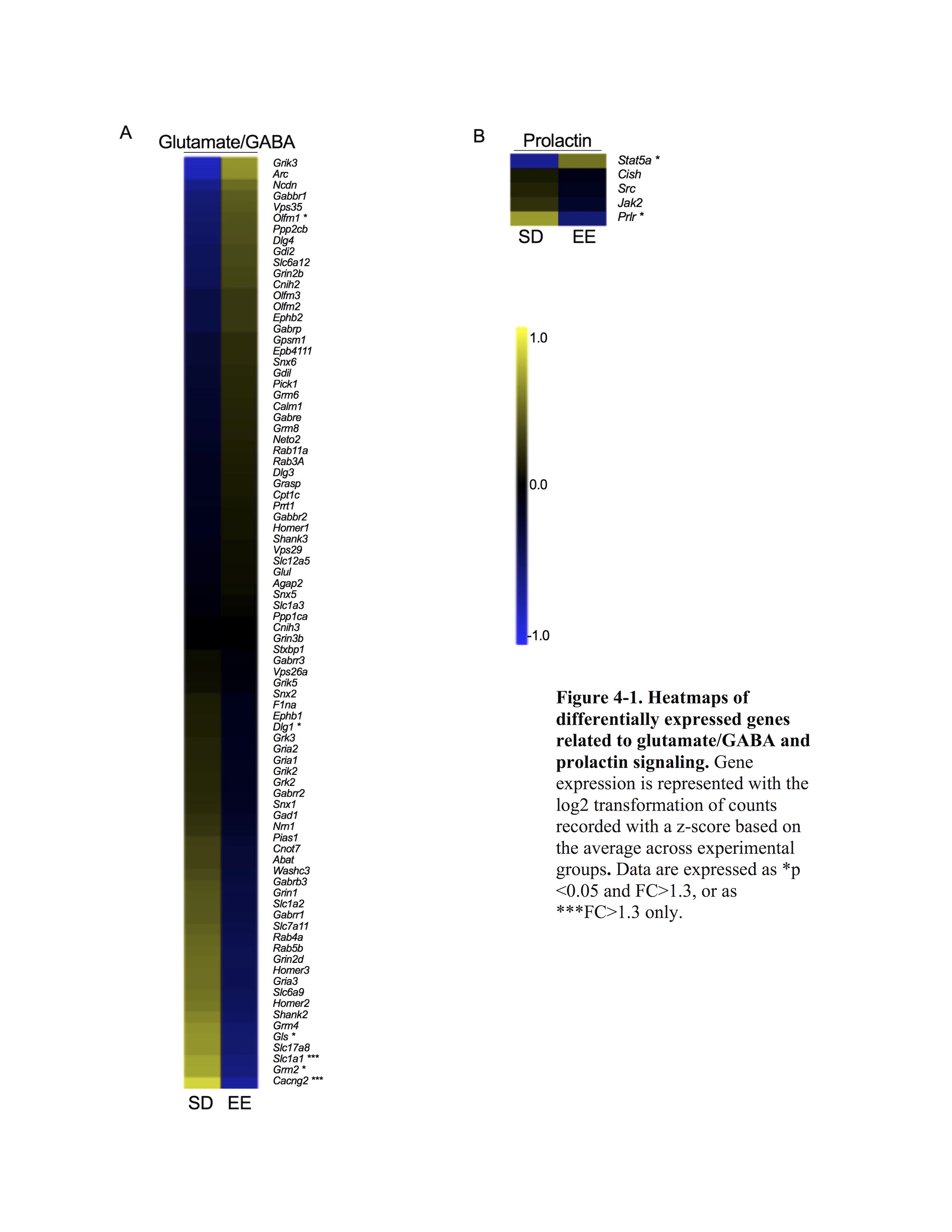

### Extended Data Figure 5-1

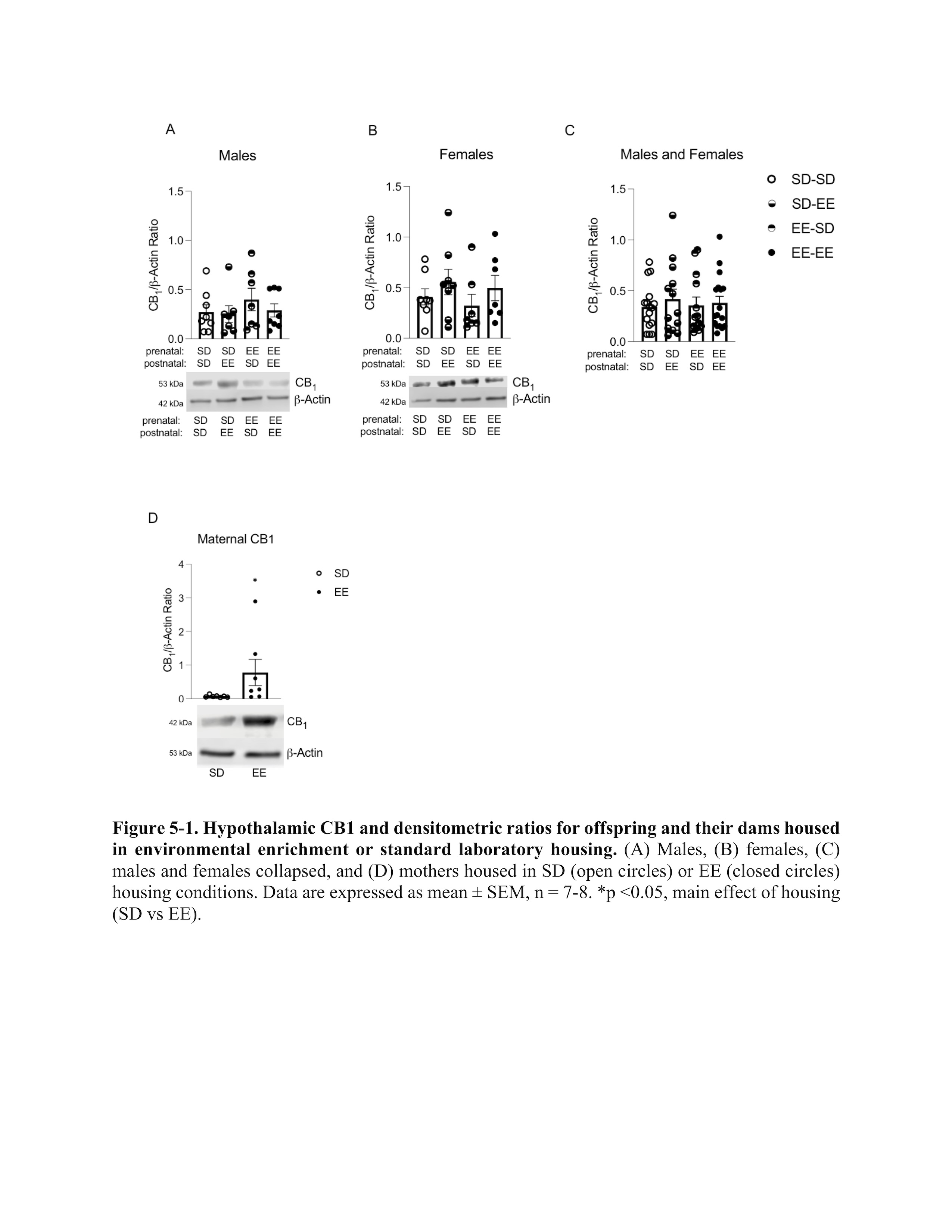
