## Extended Data Table 3-1 for "Milking it for all it’s worth: The effects of environmental enrichment on maternal nurturance, lactation quality, and offspring social behavior"

| Extended Table 3-1. Genes identified with DESeq analysis. | | | |
| --- | --- | --- | --- |
| Symbols | Log2Fold Change | p value | padj |
| Akap9 | -0.922 | 4.46E-15 | 4.23E-11 |
| LOC100361083 | -1.089 | 8.54E-10 | 4.05E-06 |
| Ada | 0.875 | 1.48E-09 | 4.68E-06 |
| Abhd2 | -0.963 | 6.12E-09 | 1.45E-05 |
| Atl2 | -0.681 | 7.64E-09 | 1.45E-05 |
| Golgb1 | -0.849 | 1.54E-08 | 2.34E-05 |
| Trip11 | -0.623 | 1.73E-08 | 2.34E-05 |
| Nop9 | 0.446 | 7.06E-08 | 8.37E-05 |
| Tjp3 | 0.436 | 1.25E-07 | 0.000131 |
| Cd14 | -1.31 | 5.86E-07 | 0.000537 |
| Dpyd | -0.47 | 6.23E-07 | 0.000537 |
| Yes1 | -0.686 | 6.85E-07 | 0.000541 |
| Atl3 | -0.529 | 1.15E-06 | 0.000839 |
| Tbc1d14 | -0.447 | 1.39E-06 | 0.000942 |
| Umps | 0.651 | 2.97E-06 | 0.00176 |
| Ccng2 | -0.815 | 4.03E-06 | 0.00213 |
| Angptl1 | -1.381 | 5.55E-06 | 0.00277 |
| Alcam | -0.931 | 6.63E-06 | 0.00314 |
| Klhl24 | -0.676 | 7.02E-06 | 0.00317 |
| Pkd1 | -0.62 | 7.78E-06 | 0.00321 |
| Abhd14a | -0.573 | 7.71E-06 | 0.00321 |
| Tubb2a | 0.669 | 9.93E-06 | 0.00393 |
| Atg13 | -0.447 | 1.04E-05 | 0.00396 |
| Zfp593 | 0.64 | 1.09E-05 | 0.00399 |
| Fam185a | 0.707 | 1.22E-05 | 0.00428 |
| Slc30a4 | -0.512 | 1.39E-05 | 0.00469 |
| Ppm1k | -0.612 | 1.58E-05 | 0.00515 |
| Lamb2 | -1.226 | 2.22E-05 | 0.00621 |
| Tnfrsf21 | -0.481 | 2.15E-05 | 0.00621 |
| Mrps2 | 0.41 | 2.23E-05 | 0.00621 |
| Mcrip2 | 0.676 | 2.44E-05 | 0.00662 |
| Gab1 | -0.406 | 2.57E-05 | 0.00668 |
| Suco | -0.819 | 0.000028 | 0.00681 |
| Dhrs7 | -0.954 | 3.11E-05 | 0.00738 |
| Mrps26 | 0.414 | 3.35E-05 | 0.00755 |
| Symbols | Log2Fold Change | p value | padj |
| Actb | 0.488 | 3.27E-05 | 0.00755 |
| Tbc1d2 | 0.601 | 3.63E-05 | 0.00782 |
| Ifitm3 | -1.238 | 3.81E-05 | 0.00799 |
| Arhgef3 | -0.663 | 3.96E-05 | 0.00799 |
| Casp9 | 0.512 | 3.89E-05 | 0.00799 |
| Fh | 0.46 | 4.15E-05 | 0.00821 |
| Pus1 | 0.416 | 4.67E-05 | 0.00898 |
| Pabpc1 | -0.506 | 5.06E-05 | 0.00942 |
| Dus2 | 0.523 | 5.27E-05 | 0.00961 |
| Angpt1 | -0.758 | 0.000055 | 0.00966 |
| Angptl2 | 1.831 | 5.44E-05 | 0.00966 |
| Gsk3b | -0.531 | 5.95E-05 | 0.0101 |
| Nr2c2 | -0.46 | 6.43E-05 | 0.0107 |
| 8-Sep | 0.726 | 6.52E-05 | 0.0107 |
| Stxbp6 | -0.464 | 7.57E-05 | 0.012 |
| Trex1 | 0.562 | 8.02E-05 | 0.0123 |
| Socs1 | 1.08 | 7.88E-05 | 0.0123 |
| Lnpep | -0.657 | 9.86E-05 | 0.0148 |
| Lifr | -0.739 | 0.000101 | 0.0149 |
| Ccdc181 | 1.03 | 0.000114 | 0.0164 |
| Mpzl2 | -0.689 | 0.000149 | 0.0208 |
| Ptk7 | -1.021 | 0.000157 | 0.021 |
| Trim14 | 0.439 | 0.000157 | 0.021 |
| Bysl | 0.442 | 0.000162 | 0.0214 |
| Mtmr12 | -0.45 | 0.000177 | 0.023 |
| Tmtc4 | -0.644 | 0.000185 | 0.0237 |
| Sft2d1 | -0.673 | 0.000194 | 0.024 |
| RGD1359127 | 0.383 | 0.00019 | 0.024 |
| Pusl1 | 0.488 | 0.000201 | 0.0244 |
| Abcc10 | -0.845 | 0.000204 | 0.0245 |
| Tlr2 | -0.749 | 0.000227 | 0.0269 |
| Dhx58 | 0.431 | 0.00023 | 0.0269 |
| Slc20a2 | -0.905 | 0.000254 | 0.0284 |
| Srebf1 | -0.688 | 0.000248 | 0.0284 |
| Ckb | 1.14 | 0.000253 | 0.0284 |
| Symbols | Log2Fold Change | p value | padj |
| Dynll1 | 0.415 | 0.000265 | 0.0289 |
| Dhx37 | 0.671 | 0.000262 | 0.0289 |
| Cadm4 | -0.903 | 0.000294 | 0.0306 |
| Tlr5 | -0.858 | 0.000289 | 0.0306 |
| Ssr4 | -0.501 | 0.000297 | 0.0306 |
| Rbfox2 | -0.465 | 0.000307 | 0.031 |
| Slc45a3 | -1.538 | 0.000316 | 0.0312 |
| Macf1 | -0.48 | 0.000316 | 0.0312 |
| Zbtb44 | -0.454 | 0.000322 | 0.0315 |
| Pfkfb3 | -1.893 | 0.00037 | 0.0325 |
| Pigl | -0.696 | 0.000366 | 0.0325 |
| Ppm1l | -0.661 | 0.000365 | 0.0325 |
| Rbm33 | -0.408 | 0.000346 | 0.0325 |
| Preb | 0.387 | 0.000357 | 0.0325 |
| Pmm2 | 0.4 | 0.000339 | 0.0325 |
| RGD1562136 | 0.607 | 0.000362 | 0.0325 |
| Rnaseh2b | 1.114 | 0.000346 | 0.0325 |
| Adcy10 | -0.681 | 0.00038 | 0.033 |
| Tlcd1 | -1.177 | 0.000412 | 0.0349 |
| Med8 | 0.508 | 0.000424 | 0.0353 |
| Apex2 | 0.546 | 0.000421 | 0.0353 |
| Rnd1 | 0.569 | 0.000437 | 0.036 |
| Lck | 0.966 | 0.000451 | 0.0363 |
| Ovca2 | 0.568 | 0.000522 | 0.0405 |
| Wdr12 | 0.489 | 0.000536 | 0.0408 |
| Mpzl1 | -0.729 | 0.000554 | 0.0414 |
| Rpia | 0.692 | 0.000554 | 0.0414 |
| Sectm1b | -0.971 | 0.000574 | 0.0419 |
| Clip1 | -0.631 | 0.00057 | 0.0419 |
| Cdt1 | 1.183 | 0.000573 | 0.0419 |
| Mefv | 0.967 | 0.000586 | 0.0424 |
| Gpat4 | -0.613 | 0.000625 | 0.0449 |
| Rab3il1 | 1.636 | 0.000658 | 0.0466 |
| Rcc1l | 0.454 | 0.00067 | 0.0471 |
| Pon2 | -0.773 | 0.000705 | 0.0475 |
| Symbols | Log2Fold Change | p value | padj |
| Lexm | 0.853 | 0.000705 | 0.0475 |
| Nat8 | 1.705 | 0.000689 | 0.0475 |
| Zfp516 | 0.413 | 0.000717 | 0.0479 |
| Glycam1 | -0.968 | 0.000743 | 0.049 |
| Amigo3 | 0.61 | 0.000744 | 0.049 |
